# Rab11-dependent recycling of calcium channels is mediated by auxiliary subunit α_2_δ-1 but not α_2_δ-3

**DOI:** 10.1101/2021.02.25.432839

**Authors:** James O. Meyer, Annette C. Dolphin

## Abstract

N-type voltage-gated calcium channels (Ca_V_2.2) are predominantly expressed at presynaptic terminals, and their function is regulated by auxiliary α_2_δ and β subunits. All four mammalian α_2_δ subunits enhance calcium currents through Ca_V_1 and Ca_V_2 channels, and this increase is attributed, in part, to increased Ca_V_ expression at the plasma membrane. In the present study we provide evidence that α_2_δ-1, like α_2_δ-2, is recycled to the plasma membrane through a Rab11a-dependent endosomal recycling pathway. Using a dominant-negative Rab11a mutant, Rab11a(S25N), we show that α_2_δ-1 increases plasma membrane Ca_V_2.2 expression by increasing the rate and extent of net forward Ca_V_2.2 trafficking in a Rab11a-dependent manner. Dominant-negative Rab11a also reduces the ability of α_2_δ-1 to increase Ca_V_2.2 expression on the cell-surface of hippocampal neurites. In contrast, α_2_δ-3 does not enhance rapid forward Ca_V_2.2 trafficking, regardless of whether Rab11a(S25N) is present. In addition, whole-cell Ca_V_2.2 currents are reduced by co-expression of Rab11a(S25N) in the presence of α_2_δ-1, but not α_2_δ-3. Taken together these data suggest that α_2_δ subtypes participate in distinct trafficking pathways which in turn influence the localisation and function of Ca_V_2.2.

**Summary statement:** The calcium channel auxiliary subunit α_2_δ-1 but not α_2_δ-3 participates in Rab11a-dependent recycling, which in turn influences the localisation and function of Ca_V_2.2.

## Introduction

The Ca_V_1 and Ca_V_2 voltage-gated calcium channels associate with auxiliary α_2_δ and β subunits (Dolphin, 2018b). The α_2_δ subunits are glycosyl-phosphatidylinositol-anchored extracellular proteins (Davies et al., 2010; De Jongh et al., 1989; Jay et al., 1991). There are four known mammalian α_2_δ isoforms (α_2_δ-1-4) which are thought to have similar functions and topological features, despite a surprising lack of sequence homology between them (Dolphin, 2016). The α_2_δ subunits have long been known to enhance whole-cell currents through Ca_V_ channels (Barclay et al., 2001; Felix et al., 1997; Gurnett et al., 1996; Wakamori et al., 1999). However, to date there is limited evidence that α_2_δs strongly influence single-channel properties of Ca_V_ channels (Wakamori et al., 1999). As such, α_2_δ-mediated enhancement of Ca_V_ currents has often been attributed primarily to increased channel trafficking and plasma membrane insertion (Canti et al., 2005; Cassidy et al., 2014; Davies et al., 2006; Hendrich et al., 2008; Wakamori et al., 1999).

The development of functional exofacially-tagged Ca_V_2 constructs (Cassidy et al., 2014), has allowed us to define populations of plasma membrane-inserted Ca_V_2 channels using imaging approaches. This method was previously used to demonstrate that heterologous expression of α_2_δ- 1 increases cell-surface Ca_V_2.2 expression, consistent with earlier models of α_2_δ-1 function (Cassidy et al., 2014). However, subsequent studies on the proteolytic processing of α_2_δ have found that unprocessed α_2_δ-1 is unable to enhance macroscopic Ca_V_ currents despite increasing cell-surface Ca_V_2.2 expression in cell lines (Ferron et al., 2018; Kadurin et al., 2016). These studies provide convincing evidence that the α_2_δ-mediated increase in Ca_V_ currents is dependent on an increase in cell-surface Ca_V_2.2 expression, but also requires a molecular activation switch, triggered by the proteolytic cleavage of α_2_δ into α_2_ and δ. Presently, it is unclear whether α_2_δ subunits differ significantly in their ability to enhance Ca_V_ currents, and it remains to be seen if α_2_δ subtypes have uniform trafficking mechanisms particularly with regard to their effect on Ca_V_ localisation.

In a previous study (Tran-Van-Minh and Dolphin, 2010), we found that the cell-surface localization of α_2_δ-2, following its heterologous expression, was increased by recycling from Rab11a-dependent endosomes back to the plasma membrane. Rab11a belongs to the Rab family of small GTPases which regulate a multitude of intracellular trafficking pathways, facilitating membrane targeting, cargo sorting and vesicle fusion events through recruitment of effectors and direct interactions with cargo proteins (Li and Marlin, 2015). Rab11 proteins are particularly involved in trafficking through recycling endosomes (Ren et al., 1998; Ullrich et al., 1996), and inhibition of Rab11a-dependent recycling can be achieved through expression of the dominant-negative mutant Rab11a(S25N), which is locked in an inactive GDP-bound conformation (Eisfeld et al., 2011). Cell-surface expression of α_2_δ-2 is reduced in the presence of Rab11a(S25N) (Tran-Van-Minh and Dolphin, 2010), and this occludes the effect of the α_2_δ ligand gabapentin, which itself reduces cell-surface expression of α_2_δ-1 and α_2_δ-2 (Hendrich et al., 2008; Tran-Van-Minh and Dolphin, 2010). These data provide a mechanistic explanation for the action of gabapentin, whereby α_2_δ bound to gabapentin is prevented from recycling to the plasma membrane via Rab11-positive endosomes, resulting in a loss of α_2_δ cell-surface expression (Tran-Van-Minh and Dolphin, 2010). Mutational studies have revealed that gabapentin acts by binding to a site including an RRR motif in the α_2_ domain, which is present in α_2_δ-1 and α_2_δ-2 (Field et al., 2006; Gee et al., 1996; Lotarski et al., 2014; Lotarski et al., 2011). However, since α_2_δ-3 is also widely distributed in the nervous system, if there were functional differences, for example between α_2_δ-1 and α_2_δ-3, this might influence the trafficking and localisation of associated Ca_V_ channels.

In the present study, we wished to determine to what extent the enhancement of plasma membrane Ca_V_2.2 expression, a previously-demonstrated feature of α_2_δ-1 function, is conserved for α_2_δ-3. We then sought to elucidate further the mechanisms through which α_2_δs increase cell-surface Ca_V_2.2, and to understand how this process influences the localization and function of Ca_V_2.2 in cell lines and hippocampal neurons. We find that α_2_δ subunits have a differential effect on Ca_V_2.2 trafficking; although all α_2_δ proteins tested increase Ca_V_2.2 cell-surface expression to varying extents, only α_2_δ-1 and α_2_δ-2 rapidly increase net forward trafficking of Ca_V_2.2; by contrast α_2_δ-3 does not influence this process, within a 45-minute time-frame. Further examination showed that the enhanced forward trafficking of Ca_V_2.2 is a Rab11a-dependent effect and that α_2_δ-1 and α_2_δ-2 participate in Rab11a-dependent recycling, whereas α_2_δ-3 does not.

## Results

### Rab11a-dependent recycling affects α_2_δ-1 but not α_2_δ-3 cell-surface expression

Trafficking through Rab11a-positive recycling endosomes enhances α_2_δ-2 membrane expression, a pathway that is inhibited by gabapentin (Tran-Van-Minh and Dolphin, 2010). The α_2_ domain of both α_2_δ-1 and α_2_δ-2 contains a triple RRR motif, the third R of which is required for gabapentin binding, and is absent from the α_2_δ-3 and α_2_δ-4 sequences (Davies et al., 2006; Wang et al., 1999). Here, we considered the possibility that α_2_δ-1 and α_2_δ-2 enhance cell-surface Ca_V_2.2 expression, at least in part, by facilitating the recycling of Ca_V_2.2 through Rab11a-positive endosomes. Further to this, we speculated that α_2_δ-3 may traffic independently of this pathway.

To test this hypothesis, we have used the dominant-negative Rab11a(S25N) mutant, which we found previously to inhibit cell-surface expression of α_2_δ-2 (Tran-Van-Minh and Dolphin, 2010), and thus a similar result was expected for α_2_δ-1. We compared cell-surface expression of HA-tagged α_2_δ-1 and α_2_δ-3, in the absence and presence of Rab11a(S25N) (Fig. 1A, B). Cell-surface α_2_δ-3 immunostaining is typically weaker than that for α_2_δ-1 (Fig. 1A), so we used antigen retrieval to maximise available HA signal. We found a reduction in cell-surface α_2_δ-1-HA staining of 47 %, when co-expressed with Rab11a(S25N) (Fig. 1B). In contrast α_2_δ-3 showed no significant difference in cell-surface expression whether expressed with or without Rab11a(S25N) (Fig. 1B). These data provide evidence that α_2_δ-1, like α_2_δ-2 (Tran-Van-Minh and Dolphin, 2010), is regulated by Rab11a-dependent recycling and suggest that cell-surface expression of α_2_δ-3 is not promoted by trafficking through this endosomal recycling pathway (Fig. 1D). This result was supported by measurement of the intracellular α_2_δ levels which were strongly reduced by Rab11a(S25N) in the case of α_2_δ-1 but not α_2_δ-3 (Fig. 1A, C). This suggests that block of recycling endosome function may lead re-routing of α_2_δ-1 to degradation pathways (Fig. 1D).

**Figure 1.**
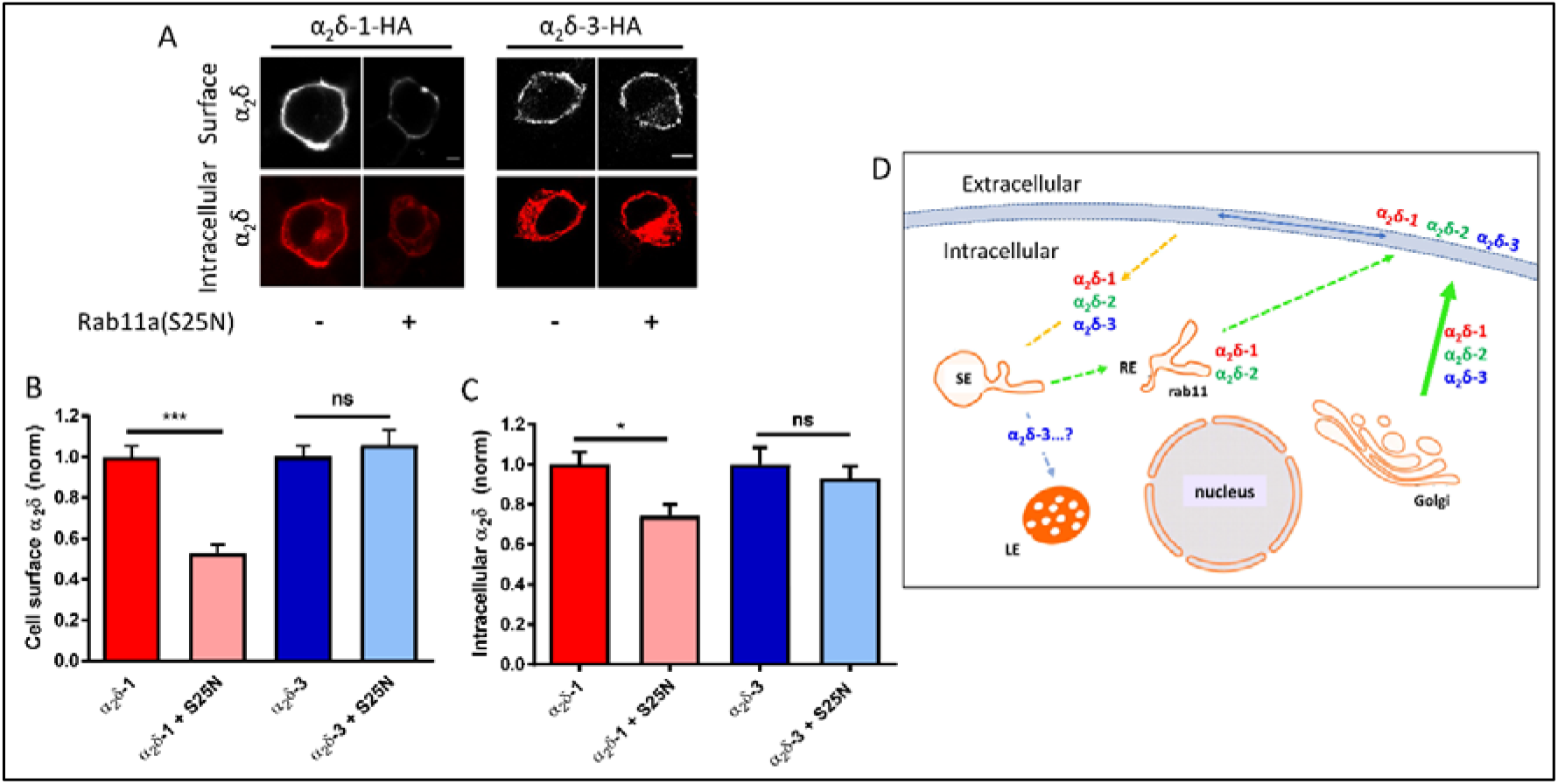
Dominant-negative Rab11a(S25N) reduces steady-state cell-surface expression of HA-tagged α_2_δ-1 but not α_2_δ-3. **(A)** Confocal images of cell-surface (top row) and intracellular (bottom row) HA staining in N2a cells expressing α_2_δ-1-HA (left) or α_2_δ-3-HA (right) in the absence or presence of Rab11a(S25N). Scale bars (5μm) refer to either α_2_δ-1 or α_2_δ-3 expressing cells. Cell surface HA was detected in non-permeabilised cells and intracellular HA subsequently detected following permeabilization (see Methods). **(B)** Normalized mean cell-surface α_2_δ expression for α_2_δ-1 control (red), α_2_δ-1 + Rab11a(S25N) (pink), α_2_δ-3 control (blue), α_2_δ-3 + Rab11a(S25N) (light blue). **(C)** Normalized mean intracellular α_2_δ expression for α_2_δ-1 control (red), α_2_δ-1 + Rab11a(S25N) (pink), α_2_δ-3 control (blue), α_2_δ-3 + Rab11a(S25N) (light blue). For (B) and (C), mean fluorescence intensity per cell was normalized to mean α_2_δ control conditions for each experiment. Normalised data were then pooled from 3 separate experiments and plotted as mean ± SEM. Each cell was measured for cell-surface and intracellular HA expression: α_2_δ-1 control (n = 159 cells), α_2_δ-1 + Rab11a(S25N) (n = 161 cells), α_2_δ-3 control (n = 134 cells), α_2_δ-3 + Rab11a(S25N) (n = 135 cells). Statistical significance was determined using Student’s unpaired t test; *, *P* < 0.05; ***, *P* < 0.001. **(D)** Diagram of pathways for α_2_δ membrane expression and recycling, and Rab11 expression. SE = sorting endosome, RE = recycling endosome, LE = late endosome/lysosome.

### Cell-surface Ca_V_2.2 expression is reduced by Rab11a(S25N) when α_2_δ-1 or α_2_δ-2 are present

In our previous study (Tran-Van-Minh and Dolphin, 2010), we found that inhibition of α_2_δ-2 recycling through Rab11a-positive endosomes reduced whole-cell Ca_V_2.2 currents. However, a direct effect on plasma membrane insertion of associated Ca_V_ channels was not examined. Here we compared the plasma membrane expression of exofacially-tagged Ca_V_2.2, containing a bungarotoxin binding site (BBS) tag (Cassidy et al., 2014), when co-expressed with β1b, either alone or together with α_2_δ- 1, or α_2_δ-3. We then examined the effect of Rab11a(S25N) (Fig. 2A). Plasma membrane-inserted Ca_V_2.2 was increased by 92 % and 38 % when co-expressed with α_2_δ-1 and α_2_δ-3, respectively (Fig. 2B), in agreement with our previous study (Kadurin et al., 2016). However, Rab11a(S25N) co-expression reduced the plasma membrane level of Ca_V_2.2 by 53 % when α_2_δ-1 was present, but had no effect in the presence of α_2_δ-3, or in the absence of α_2_δ (Fig. 2B). Total Ca_V_2.2 levels were also reduced by Rab11a(S25N) in the presence of α_2_δ-1 but not in the absence of α_2_δ or presence of α_2_δ- 3 (Fig. 2A, C). Following these findings, we also compared the effect of Rab11a(S25N) on plasma membrane Ca_V_2.2 in the presence α_2_δ-2 (Fig. S1A, B). As expected, cell-surface Ca_V_2.2 expressed with α_2_δ-2 and β1b was reduced by 35 % by Rab11a(S25N) (Fig. S1B).

**Figure 2.**
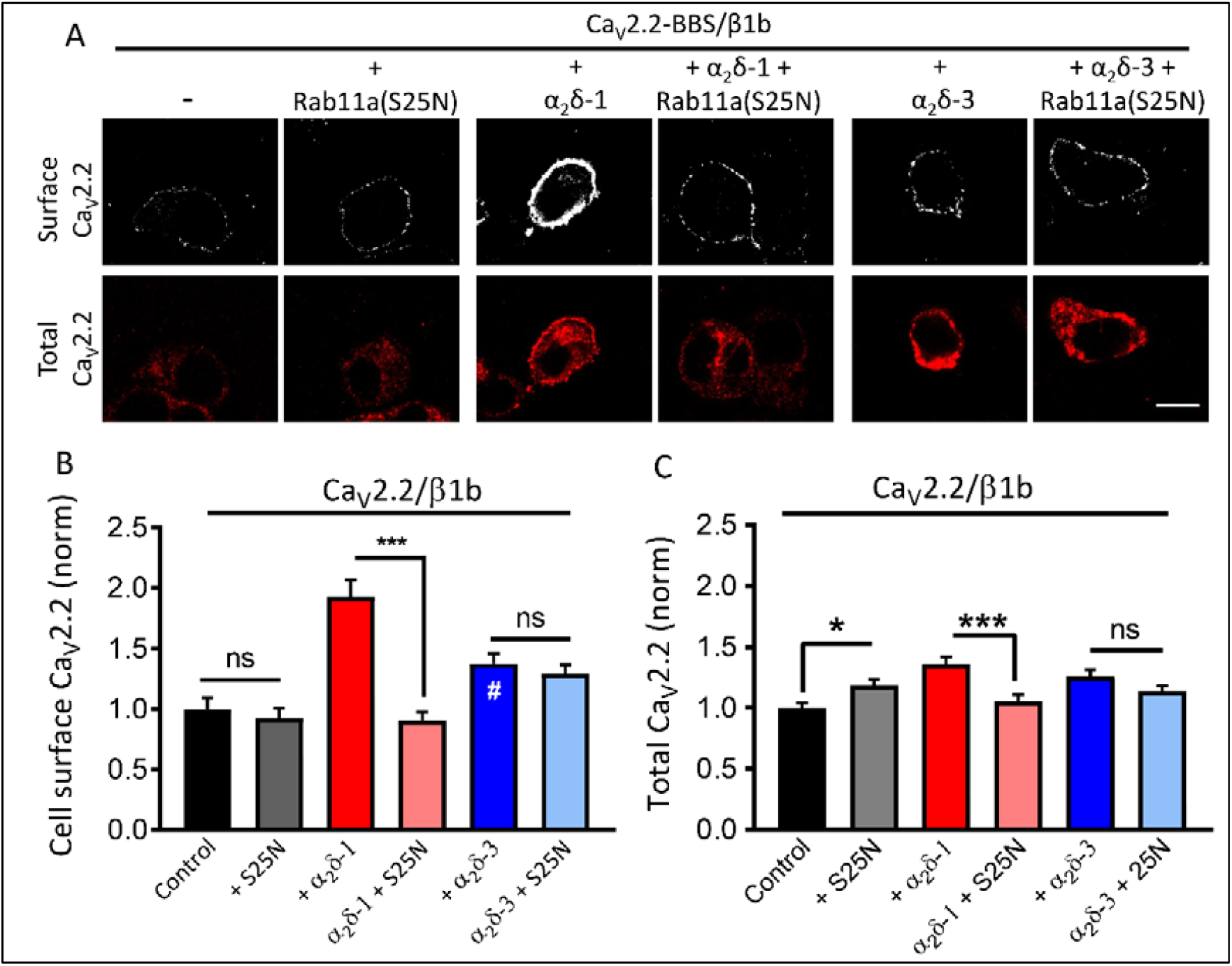
Steady-state cell-surface Ca_V_2.2 is reduced by Rab11a(S25N) when expressed with α_2_δ-1, but not α_2_δ-3. **(A)** Confocal images of cell-surface (top row, BBS) and total (bottom row, II-III loop Ab) Ca_V_2.2-BBS expressed in N2a cells with β1b and either: no α_2_δ, α_2_δ-1, or α_2_δ-3, in the absence or presence of Rab11a(S25N). Scale bar applies to all images = 5μm. **(B)** Mean cell-surface Ca_V_2.2-BBS levels expressed in N2a cells with β1b and: no α_2_δ (black, n = 142), no α_2_δ + Rab11a(S25N) (grey, n = 145 cells), α_2_δ-1 (red, n = 148 cells), α_2_δ-1 + Rab11a(S25N) (pink, n = 140 cells), α_2_δ-3 (blue, n = 132 cells), α_2_δ-3 + Rab11a(S25N) (light blue, n = 144 cells), normalized to the control level in the absence of α_2_δ. Statistical significance was determined using one-way ANOVA and Bonferroni post hoc tests; ns, *P* > 0.05; ***, *P* < 0.001; #, *P* < 0.05 vs no α_2_δ. **(C)** Total Ca_V_2.2-BBS in cells shown in (B). Statistical significance was determined using one-way ANOVA and Bonferroni post hoc tests; ns, *P* > 0.05; *, *P* < 0.05; ***, *P* < 0.001. For (B) and (C), each cell was measured for cell-surface BBS expression in non-permeabilised cells using α-BTX-AF488, and for total Ca_V_2.2 following permeabilization using the II-III loop Ab (see Methods). Fluorescence intensity per cell was normalized to mean Ca_V_2.2/β1b control condition for each experiment. Normalized data were then pooled from 3 separate experiments and plotted ± SEM.

Together, these data suggest that α_2_δ-1 and α_2_δ-2 enhance cell-surface Ca_V_2.2 expression, in part through Rab11a-dependent recycling to the plasma membrane, which Ca_V_2.2 is unable to access independently of these α_2_δ subunits. In contrast, although α_2_δ-3 produced a 38 % increase in Ca_V_2.2 cell-surface expression (Fig. 2B), this was independent of a Rab11a-dependent recycling pathway, suggesting that α_2_δ-3 is only able to promote cell-surface Ca_V_2.2 trafficking through another pathway, such as direct trafficking from the Golgi apparatus.

### Rab11a(S25N) reduces Ca_V_2.2 expression in neurites of cultured hippocampal neurons when co-expressed with α_2_δ-1 but not α_2_δ-3

We have previously demonstrated that Ca_V_β subunits are able to increase plasma membrane expression of Ca_V_2.2 in non-neuronal cells, even in the absence of α_2_δ, although there is generally an approximately 2-fold increase in the presence of α_2_δ-1 (Cassidy et al., 2014; Kadurin et al., 2016) (see Fig. 2B). However, in primary hippocampal neuronal cultures, we have found the presence of α_2_δ to be critical for localisation of Ca_V_2.2 to neurites (Kadurin et al., 2016). Here, we examined whether Ca_V_2.2 expression on the surface of hippocampal neurites is also influenced by Rab11a in an α_2_δ-dependent manner. To do this we used an exofacially HA-tagged and N-terminally GFP-tagged Ca_V_2.2 construct (Macabuag and Dolphin, 2015), which allowed us to quantify cell-surface and total Ca_V_2.2 expression in non-permeabilised neurons. We expressed GFP-Ca_V_2.2-HA plus β1b, with or without α_2_δ-1 or α_2_δ-3, and with or without Rab11a(S25N) (Fig. 3A). In accordance with our previous studies (Kadurin et al., 2016), expression of Ca_V_2.2 on the cell-surface of neurites was extremely low in the absence of α_2_δ (Fig. 3A, B). Here we found a ∼4-fold increase in cell-surface Ca_V_2.2-HA expression with α_2_δ-1, and ∼3.5-fold increase with α_2_δ-3, relative to control (Fig. 3B). Expression of GFP in the neurites, representing total Ca_V_2.2, was also elevated by co-expression of both α_2_δ-1 and α_2_δ-3, by 2.1-fold and 1.9-fold, respectively (Fig. 3C). Consistent with our findings in non-neuronal cells, cell-surface Ca_V_2.2-HA was reduced by co-expression of Rab11a(S25N) in the presence of α_2_δ-1 (by ∼44 %), but was not affected in the presence of α_2_δ-3 (Fig. 3B). However, we saw no change in total Ca_V_2.2-GFP expression in the neurites with Rab11a(S25N), either in the absence or presence of α_2_δ-1 or α_2_δ-3 (Fig. 3C). Together these data suggest that Rab11a-dependent recycling enhances neurite Ca_V_2.2 cell-surface expression in an α_2_δ-selective manner.

**Figure 3.**
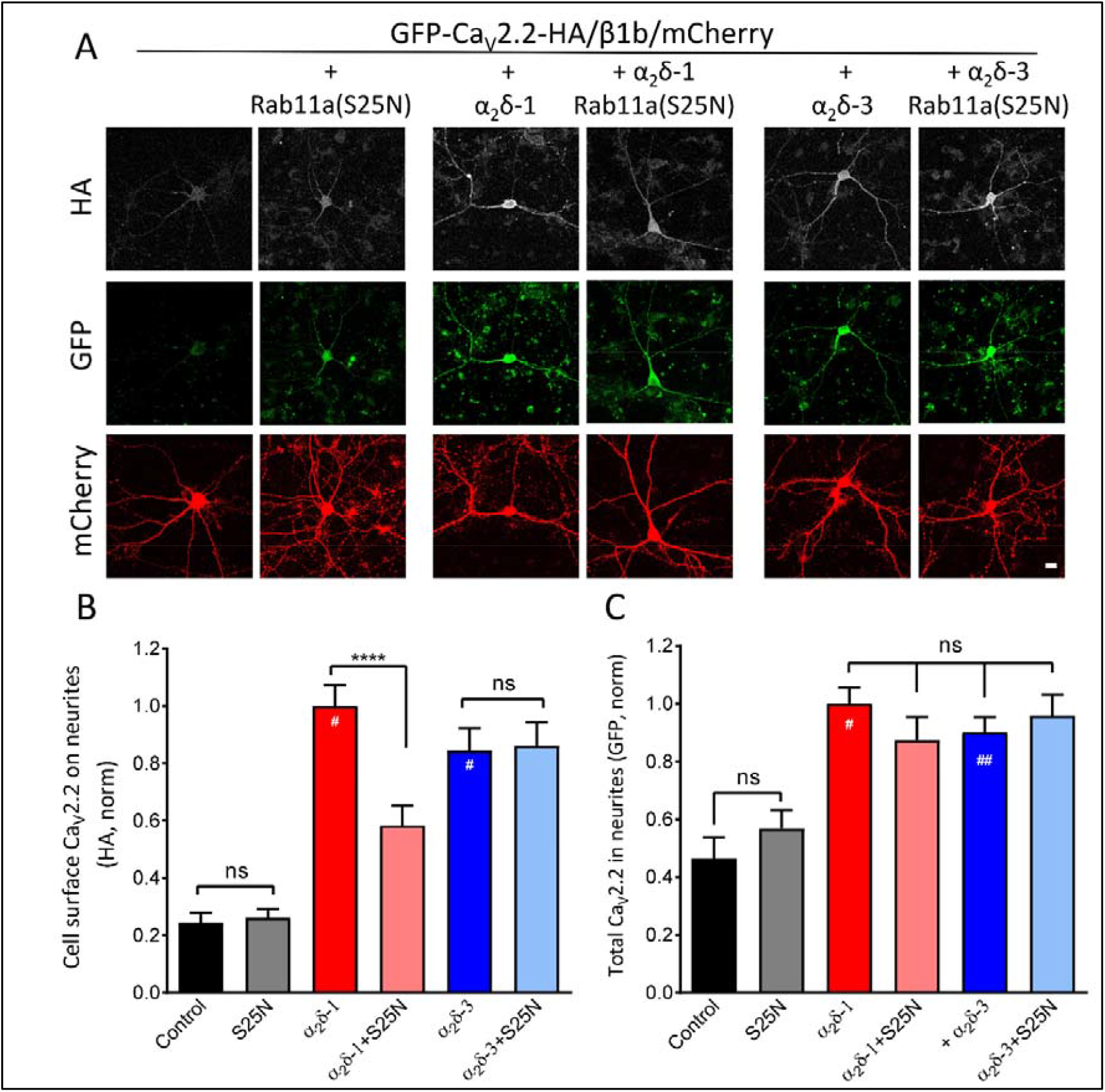
Rab11a(S25N) reduces plasma membrane-inserted Ca_V_2.2 in hippocampal neurites when expressed with α_2_δ-1 but not α_2_δ-3. **(A)** Confocal images of rat hippocampal pyramidal neurons expressing GFP-Ca_V_2.2-HA/β1b/mCherry control, + Rab11a(S25N), α_2_δ-1, α_2_δ-1 + Rab11a(S25N), α_2_δ-3 or α_2_δ-3 + Rab11a(S25N). Scale bar applies to all images = 20 μm. Top row, HA immunostaining in non-permeabilised neurons; middle row, GFP to determine total Ca_V_2.2 expression; bottom row, mCherry (transfection marker) expression. **(B)** Normalized mean cell-surface GFP-Ca_V_2.2-HA, determined from HA staining in non-permeabilised neurons, in the processes of pyramidal rat neurons expressed with β1b/mCherry for control (black), Rab11a(S25N) (grey), α_2_δ-1 (red), α_2_δ-1 + Rab11a(S25N) (pink), α_2_δ-3 (blue) or α_2_δ-3 + Rab11a(S25N) (light blue). **(C)** Normalized mean total GFP-Ca_V_2.2-HA (determined from GFP density) in the processes of pyramidal rat neurons expressed with β1b/mCherry for control (black), Rab11a(S25N) (grey), α_2_δ-1 (red), α_2_δ-1 + Rab11a(S25N) (pink), α_2_δ-3 (blue) or α_2_δ-3 + Rab11a(S25N) (light blue). For (B) and (C), between 2-5 neuronal processes were measured and averaged per cell, 10-15 cells per condition per experiment, for a total of 3 transfections. Total cell numbers for each condition: control (n = 30 cells), Rab11a(S25N) (n = 19 cells), α_2_δ-1 (n = 51 cells), α_2_δ-1 + Rab11a(S25N) (n = 42 cells), α_2_δ-3 (n = 46) and α_2_δ-3 + Rab11a(S25N) (n = 40 cells). Normalized mean fluorescence intensity per neuron was pooled from three separate experiments and data are plotted ± SEM values. Statistical significance was determined using one-way ANOVA and Bonferroni post hoc tests; ns, *P* > 0.05; ****, *P* < 0.0001. Significance vs no α_2_δ control: #, *P* < 0.0001; ##, *P* = 0.002.

### Rab11a(S25N) reduces whole-cell Ca_V_2.2 currents when co-expressed with α_2_δ-1 but not α_2_δ-3

Here we investigated whether the reduction in cell-surface Ca_V_2.2 expression by Rab11a(S25N) correlates to a change in whole-cell Ca_V_ currents and whether any change we observe is α_2_δ-dependent. We used patch-clamp recording to measure whole-cell Ba^2+^ currents in tsA201 cells expressing Ca 2.2/β1b/α_2_δ, with or without Rab11a(S25N) (Fig. 4A). We found that the peak Ba2+ current density was reduced by 59 % at +10 mV, when α_2_δ-1 and Rab11a(S25N) were co-expressed, compared to α_2_δ-1 expressed alone (Fig. 4A, B). Corresponding maximum conductance (G_max_) values were reduced from 1.4 ± 0.2 nS/pF to 0.7 ± 0.1 nS/pF by the presence of Rab11a(S25N) (Fig. 4C). We saw no significant changes to steady-state inactivation under the two conditions (Fig. S2A), supporting the view that current density changes occurred as a result of reduced plasma membrane Ca_V_2.2 expression rather than changes to biophysical properties of the channels.

**Figure 4.**
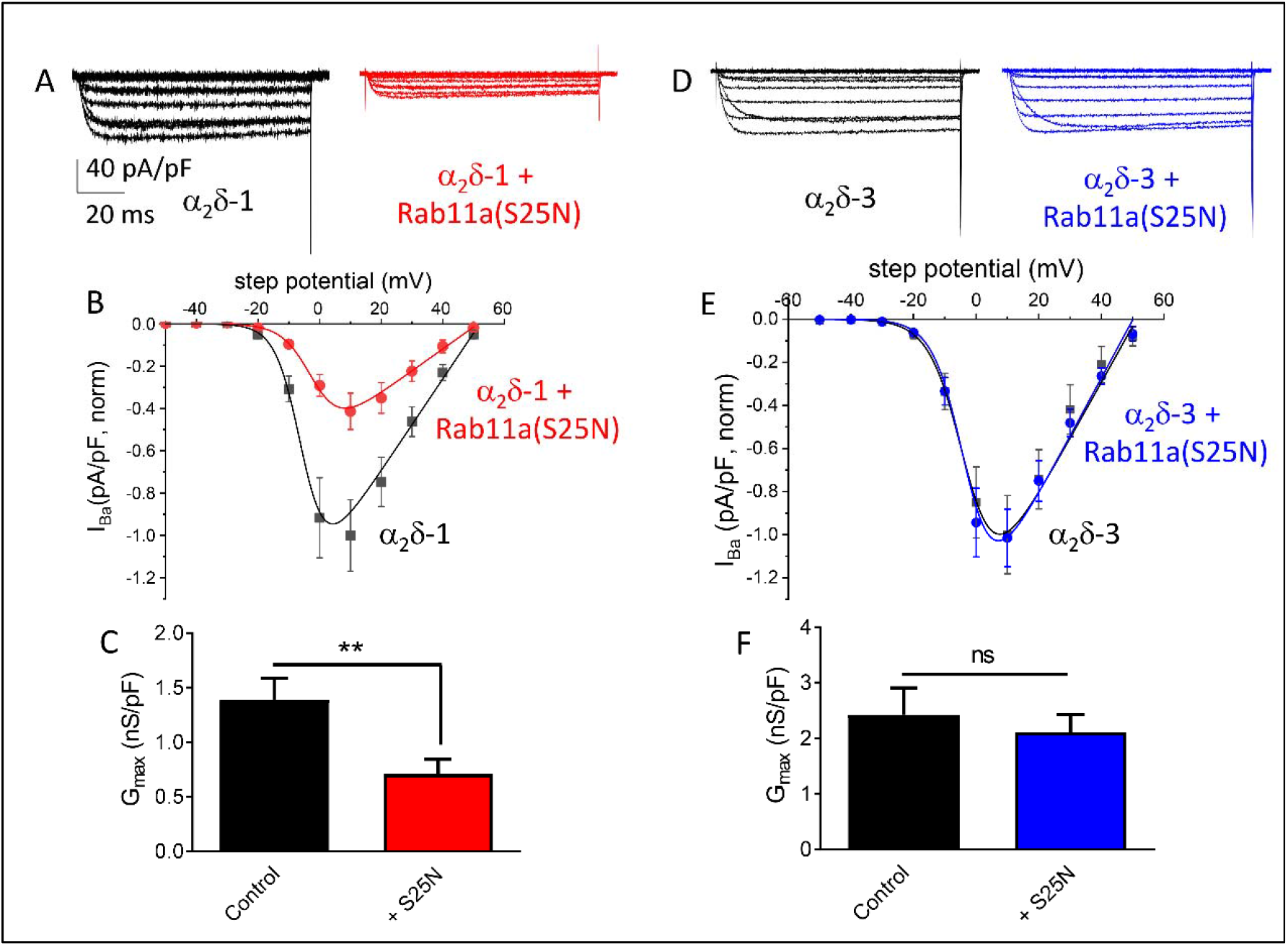
Rab11a(S25N) reduces whole-cell Ca_V_2.2 currents when expressed with α_2_δ-1, but not α_2_δ-3. **(A)** Example whole-cell current traces for Ca_V_2.2 expressed in tsA-201 cells with: β1b/α_2_δ-1 (black) and β1b/α_2_δ-1 + Rab11a(S25N) (red). Scale bars (40 pA/pF and 20 ms) refer to all traces. **(B)** Mean *IV* plots for Ca_V_2.2 with: β1b/α_2_δ-1 (black squares, n = 15) and β1b/α_2_δ-1 + Rab11a(S25N) (red circles, n = 14), fitted by a modified Boltzmann function. The potentials for 50 % activation were −4.23 ± 0.76 mV, and −1.39 ± 1.26 mV, respectively. **(C)** Mean *G*_max_ values obtained by fitting each individual trace in (B) to a modified Boltzmann function Ca_V_2.2/β1b/α_2_δ-1 (black), Ca_V_2.2/β1b/α_2_δ-1 + Rab11a(S25N) (red). **(D)** Example whole-cell current traces for Ca_V_2.2 with: β1b/α_2_δ-3 (black) and β1b/α_2_δ-3 + Rab11a(S25N) (blue). **(E)** Mean *IV* plots for Ca_V_2.2 with: β1b/α_2_δ-3 (black squares, n = 15) and β1b/α_2_δ-3 + Rab11a(S25N) (blue circles, n = 14), fitted by a modified Boltzmann function. The potentials for 50 % activation were −4.21 ± 1.59 mV and −3.93 ± 0.98 mV, respectively **(F)** Mean *G*_max_ values obtained by fitting each individual trace in (E) to a modified Boltzmann function Ca_V_2.2/β1b/α_2_δ-3 (black), Ca_V_2.2/β1b/α_2_δ-3 + Rab11a(S25N) (blue). Data are plotted as mean ± SEM. Statistical significance determined by Student’s unpaired t-test; ns, *P* > 0.05; **, *P* < 0.01.

Having established that Rab11a(S25N) reduced both α_2_δ-1-mediated Ca_V_2.2 plasma membrane expression and Ca_V_2.2 current enhancement to similar extents, it was necessary to determine whether Rab11a(S25N) would produce a reduction in α_2_δ-3-mediated Ca_V_2.2 current enhancement. In contrast with the effect of α_2_δ-1, we found no difference in mean current density (Fig. 4D, E), G_max_ (Fig. 4F)) or steady-state inactivation (Fig. S2B) when Rab11a(S25N) was co-expressed with α_2_δ-3, compared to α_2_δ-3 alone. This result is consistent with our observation that cell-surface expression of Ca_V_2.2 is unaffected by Rab11a-dependent recycling when expressed with α_2_δ-3.

### Net forward trafficking of Ca_V_2.2 is enhanced by α_2_δ-1 and α_2_δ-2, but not α_2_δ-3

Having provided evidence that α_2_δ-3 enhances steady-state plasma membrane Ca_V_2.2 expression, albeit to a smaller degree than α_2_δ-1 (Fig. 2B), we considered two possible explanations for this effect: (1) α_2_δ subunits increase forward trafficking of Ca_V_2.2 to the cell-surface, or (2) α_2_δ subunits reduce the rate of Ca_V_2.2 endocytosis from the cell-surface. Previously, we found that the rate of Ca_V_2.2 endocytosis was not altered by expression of α_2_δ-1 (Cassidy et al., 2014; Dahimene et al., 2018). In the present study, we used an α-bungarotoxin (BTX) live-labelling approach to compare the rates of net forward trafficking of Ca_V_2.2-BBS expressed with β1b either alone or together with α_2_δ-1, α_2_δ-2 or α_2_δ-3 (Fig. 5A). When compared to control conditions, we found that co-expression of α_2_δ-1 and α_2_δ-2 produced significantly higher insertion of Ca_V_2.2 into the plasma membrane, at each time point tested, starting at 15 min (Fig. 5B). In contrast, the appearance of Ca_V_2.2 on the cell-surface did not differ significantly between the control and α_2_δ-3 conditions up to 45 min, but there was a statistically significant increase of 23 % at 60 min (Fig. 5B), which is consistent with our observation of its effect on steady-state Ca_V_2.2 expression (Fig. 2B). Estimates for initial rates of net forward Ca_V_2.2 trafficking were made using the slope value of a straight line between 0-15 minutes. While there was a 2.7-fold increase in initial rate of Ca_V_2.2 appearance on the cell-surface with α_2_δ-1 co-expression compared to control (Fig. 5C), we found no significant difference in initial Ca_V_2.2 trafficking rates for the α_2_δ-3 condition. Together these data support a role for α_2_δ-1 in enhancing rapid plasma membrane-insertion of Ca_V_2.2 by increasing the rate of forward trafficking, whereas α_2_δ-3 had no clear effect on the initial rate of forward Ca_V_2.2 trafficking (Fig. 5C).

**Figure 5.**
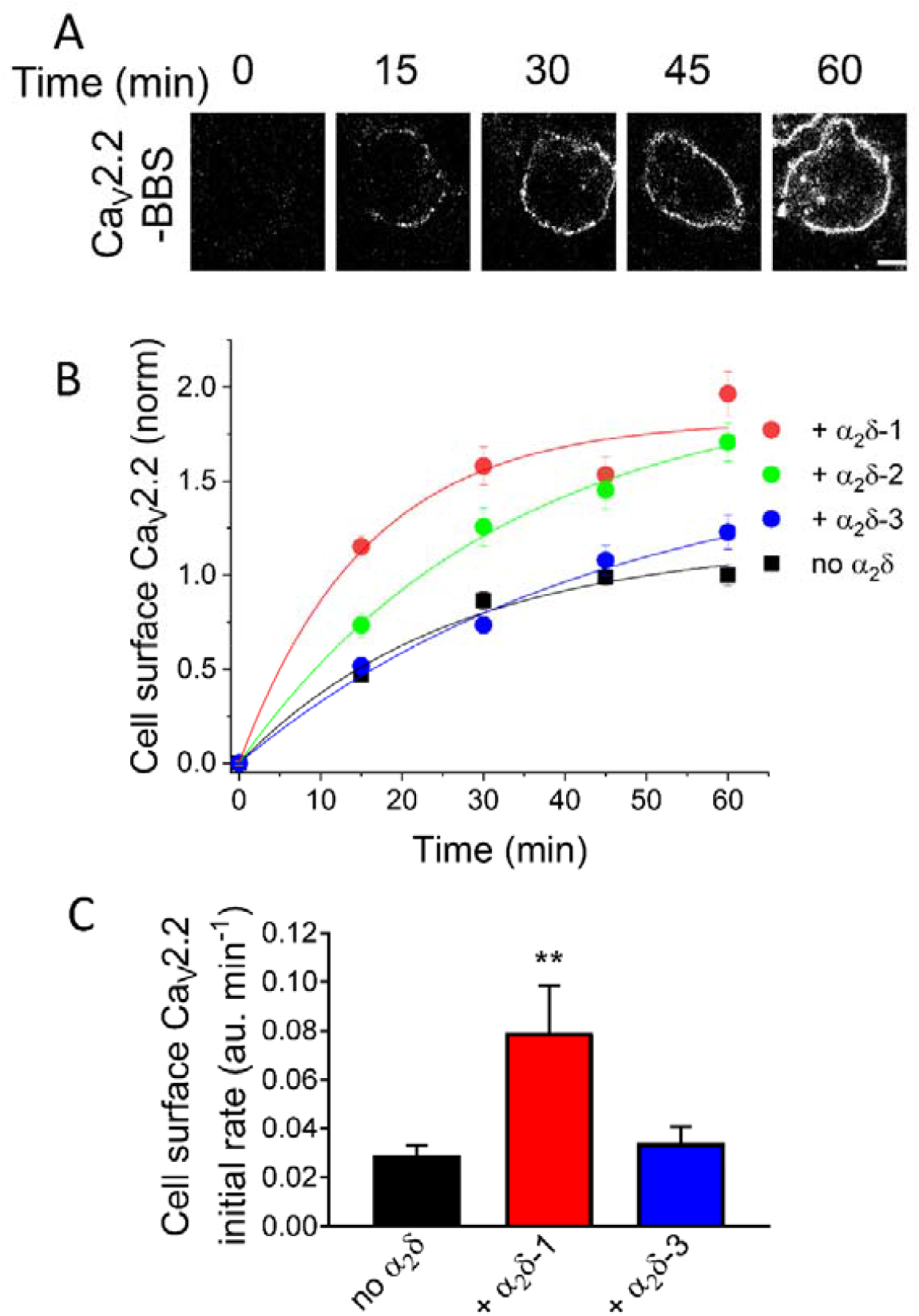
Net forward trafficking of Ca_V_2.2-BBS is increased by α_2_δ-1 and α_2_δ-2, but not by α_2_δ-3. **(A)** Example images from forward trafficking assay for Ca_V_2.2-BBS expressed in N2a cells with β1b and α_2_δ-1. Cells were live-labelled with α-BTX-AF488 for 0, 15, 30, 45, and 60 min. following pre-incubation with unlabelled α-BTX at 17ºC for 30 min. Scale bar = 5μm. **(B)** Normalized mean cell-surface Ca_V_2.2-BBS fluorescence in cells expressing Ca_V_2.2-BBS and β1b alone (black) or with either α_2_δ-1 (red), α_2_δ-2 (green) or α_2_δ-3 (blue). An average of 30 - 60 cells were analysed per time point, for each condition in an individual experiment. The data were collected as paired experiments between Ca_V_2.2/β1b (control) and Ca_V_2.2/β1b + α_2_δ (n = 3 each), with a total of 9 for Ca_V_2.2/β1b. Individual experiments were normalized to mean 60 min fluorescence of the paired Ca_V_2.2/β1b condition, before being pooled together. Data were fitted with an exponential association equation; the time constant (τ) = 26.1 min (no α_2_δ), 15.6 min (+ α_2_δ- 1), 32.4 min (+ α_2_δ-2) and 44.9 min (+ α_2_δ-3) and asymptotic normalized cell-surface Ca_V_2.2 (amplitude) = 1.17 (no α_2_δ), 1.82 (+ α_2_δ-1), 2.0 (+ α_2_δ-2) and 1.63 (+ α_2_δ-3). Statistical significance of mean fluorescence values compared to the control condition at each time point was determined using one-way ANOVA and Bonferroni post-hoc tests. For α_2_δ-1, *P* < 0.001 at all time-points; for α_2_δ- 2, *P* < 0.01 at 15 and 30 min, and *P* < 0.001 at 45 and 60 min; for α_2_δ-3, *P* < 0.05 at 60 min. **(C)** Initial rate of net forward Ca_V_2.2-BBS trafficking. The gradient of straight line between 0-15 min was obtained as an average for each individual experiment and summarised (in a.u. / min), for Ca_V_2.2/β1b control (black, n = 9), Ca_V_2.2/β1b + α_2_δ-1 (red, n = 3), and Ca_V_2.2/β1b + α_2_δ-3 (blue, n=3). Data are mean ± SEM. ** *P* = 0.0021 (one-way ANOVA and Dunnett’s multiple comparison test vs no α_2_δ).

### Dominant-negative Rab11a reduces net forward trafficking of α_2_δ-1

Next, we compared net forward trafficking rates of α_2_δ-1-BBS in the presence or absence of Rab11a(S25N) (Fig. 6A). We found that net forward trafficking of α_2_δ-1-BBS was reduced by co-expression of Rab11a(S25N), with a 41 % reduction in cell-surface α_2_δ-1-BBS after 30 minutes (Fig. 6B), consistent with the reduction in cell-surface α_2_δ-1-HA induced by Rab11a(S25N) under steady-state conditions (Fig. 1B). We then compared net forward trafficking of Ca_V_2.2-BBS, when expressed with or without α_2_δ-1 and with or without Rab11a(S25N), at two time points, 10 and 30 min (Fig. 6C - E). As expected, α_2_δ-1 enhanced net-forward trafficking of Ca_V_2.2, and Rab11a(S25N) abolished this increase at both time points. In sum, these data support the conclusion that α_2_δ-1 membrane expression is enhanced by forward-trafficking from Rab11a-positive recycling endosomes, and that Ca_V_2.2 can traffic through this pathway when co-expressed with Rab11a-sensitive α_2_δ-1, but not with α_2_δ-3 (Fig. 1D).

**Figure 6.**
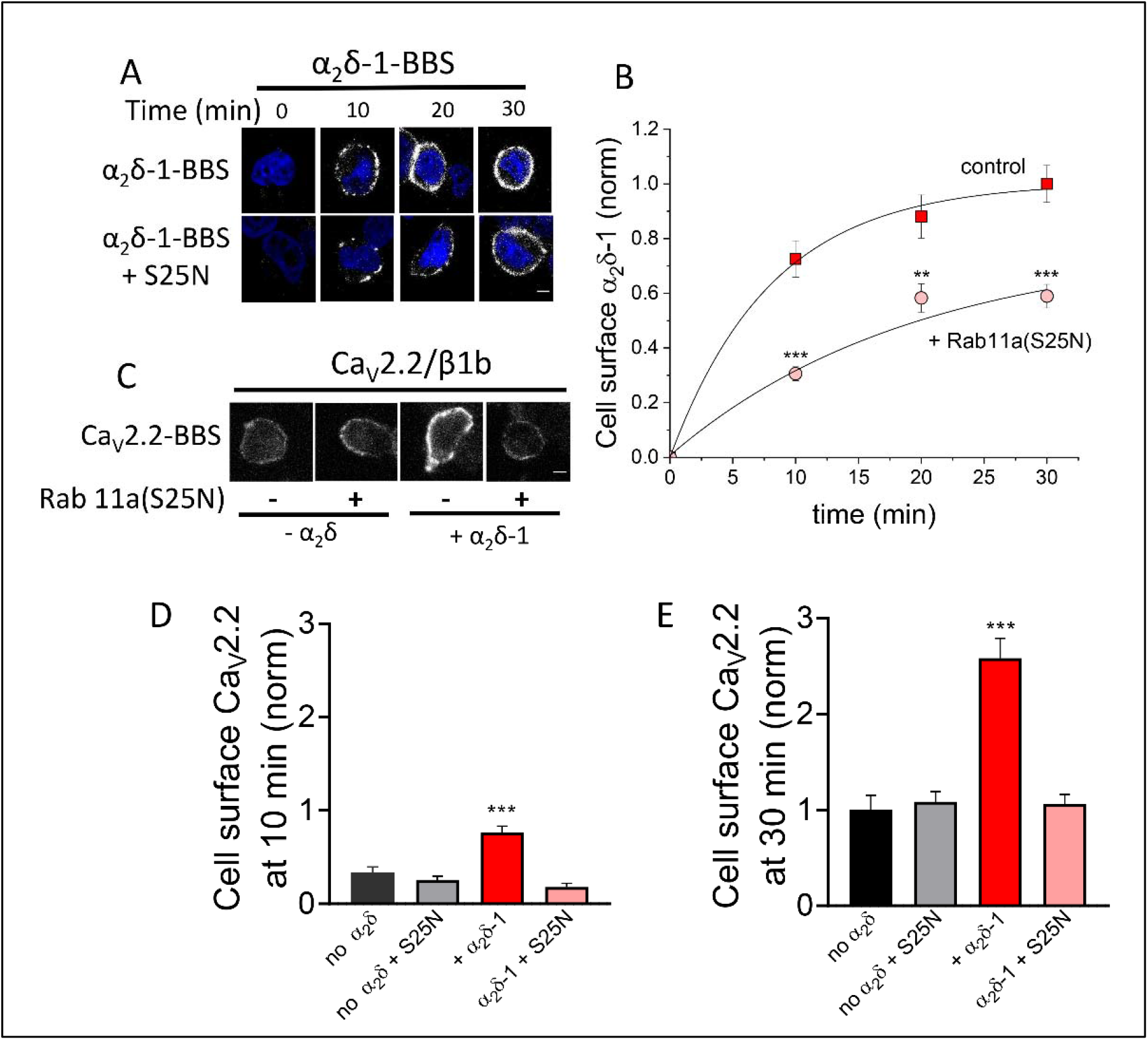
Rab11a(S25N) reduces net forward trafficking of α_2_δ-1 and Ca_V_2.2/α_2_δ-1. **(A)** Confocal images of α_2_δ-1-BBS expressed in tsA-201 cells, alone (top row) or with Rab11a(S25N) (bottom row), at each time point of net forward trafficking assay. Cells were live-labelled with α-BTX-AF488 for 0, 10, 20, and 30 min. Nuclei staining with DAPI (blue). Scale bar = 5μm. **(B)** Net forward α_2_δ-1-BBS trafficking, as in (A). Normalized mean cell-surface α_2_δ-1-BBS expressed either alone (red squares) or with Rab11a(S25N) (pink circles). Data were collected from 3 transfections with approximately 30 to 50 cells analysed per time point in each experiment. Individual experiments were normalized to the α_2_δ-1-BBS 30 min time point, before being pooled together. Data were fitted to a single exponential association, with a time constant, τ = 8.09 min for α_2_δ-1 control and 19.45 min for α_2_δ-1 + Rab11a(S25N). Statistical significance was determined using Student’s t test to compare mean fluorescence values between conditions at each time point, \*\**P*<0.01, \*\*\**P*<0.001. **(C)** Confocal images of cell-surface Ca_V_2.2-BBS expressed in N2a cells with β1b either alone or with α_2_δ-1, in the absence or presence of Rab11a(S25N) after 30 min of net forward trafficking assay. Scale bar = 5μm. **(D**,**E)** Normalized mean cell-surface Ca_V_2.2-BBS expressed in N2a cells with β1b and: no α_2_δ (black), no α_2_δ + Rab11a(S25N) (grey), + α_2_δ-1 (red) and + α_2_δ-1 + Rab11a(S25N) (pink). Cells were live-labelled with α-BTX-AF488 for 10 min (D) or 30 min (E). Fluorescence intensity per cell was normalized to mean Ca_V_2.2/β1b controls at 30 min before being pooled from 3 separate experiments and plotted as mean fluorescence ± SEM. Statistical significance was determined using one-way ANOVA and Bonferroni post hoc tests (\*\*\**P*<0.001). For (D) no α_2_δ (black, n = 105 cells), no α_2_δ + Rab11a(S25N) (grey, n = 98 cells), + α_2_δ-1 (red, n = 123 cells) and + α_2_δ-1 + Rab11a(S25N) (pink, n = 112 cells). For (E) no α_2_δ (black, n = 105 cells), no α_2_δ + Rab11a(S25N) (grey, n = 104 cells), + α_2_δ- 1 (red, n = 132 cells) and + α_2_δ-1 + Rab11a(S25N) (pink, n = 118 cells).

## Discussion

In this study we have shown that α_2_δ subunits differentially enhance plasma membrane expression of Ca_V_2.2 in both cell lines and primary neuronal cultures, and have implicated Rab11a-dependent recycling in this process. Rab11 proteins are particularly involved in trafficking through recycling endosomes (Ren et al., 1998; Ullrich et al., 1996), and here we inhibited this route through expression of the dominant-negative mutant Rab11a(S25N) (Eisfeld et al., 2011). Our data indicate that α_2_δ-1 and α_2_δ-2 promote both the steady-state cell-surface Ca_V_2.2 level and net forward trafficking of Ca_V_2.2 through a Rab11a-dependent mechanism (Fig. 1D). Blockade of this pathway leads to a reduction in cell-surface Ca_V_2.2 and α_2_δ-1 and may lead to increase in degradation, as judged by the reduction in their intracellular levels. However, α_2_δ-3 does not appear to participate in the Rab11a-dependent recycling pathway, and does not increase net forward Ca_V_2.2 trafficking in the short-term (Fig. 5B, C). Despite this, α_2_δ-3 consistently increases plasma membrane-inserted Ca_V_2.2 under steady-state conditions (Kadurin et al., 2016; and the present study), although to a smaller extent than α_2_δ-1. Inhibition of Rab11a-dependent recycling also reduces whole-cell Ca_V_2.2 currents, when expressed with α_2_δ-1, but has no effect with α_2_δ-3. Our results indicate that one of the main drivers of α_2_δ-1-mediated enhancement of Ca_V_2.2 plasma membrane-insertion is recycling back to the plasma membrane via recycling endosomes. In primary hippocampal neurites, we find that Ca_V_2.2 membrane expression is largely dependent on α_2_δ, and inhibition of Rab11a-dependent recycling also lowers cell-surface Ca_V_2.2 in neurites in the presence of α_2_δ-1.

Earlier work from this laboratory has shown that α_2_δ-2 participates in recycling through Rab11a-positive recycling endosomes and that this pathway is interrupted by gabapentin (Tran-Van-Minh and Dolphin, 2010). The gabapentin-binding RRR motif is positioned just upstream of the VWA domain in both α_2_δ-1 and α_2_δ-2, and it has been hypothesised that gabapentinoids displace an endogenous ligand in the binding pocket that includes the RRR motif (Brown et al., 1998; Field et al., 2006; Hendrich et al., 2008). Previous studies using the α_2_δ-1(R217A) or α_2_δ-2(R282A) mutants, which show much lower affinity for binding gabapentin, found them to have lower ability to enhance Ca_V_2 currents compared to their wild-type counterparts, as well as reduced endosomal recycling (Field et al., 2006; Hendrich et al., 2008; Tran-Van-Minh and Dolphin, 2010). These findings are consistent with the hypothesis that binding of an endogenous ligand in the gabapentin binding site is important for association of α_2_δ-1 and α_2_δ-2 with endosomal sorting partners. As α_2_δ-3 lacks a gabapentin binding site, we hypothesised that it would not participate in Rab11a-dependent recycling and would thereby be unable to promote forward trafficking of Ca_V_2.2 through this pathway, although the direct route via the Golgi apparatus would still be available (Tran-Van-Minh and Dolphin, 2010). In agreement with this, we find inhibition of Rab11a-dependent recycling did not influence α_2_δ-3 trafficking, unlike for α_2_δ-1 or α_2_δ-2. Presently it is unclear why α_2_δ-3 does not participate in Rab11a-dependent recycling, although α_2_δ-3 shares only 25.7 % sequence homology with α_2_δ-1 (ClustalO), which may result in conformational differences that preclude an interaction with a Rab11a effector.

Since Rab11a is a membrane-associated cytoplasmic protein, and α_2_δ-1 is extracellular and therefore present in the lumen of recycling endosomes, any functional interaction between the two will involve other proteins. There are several studies that have examined the interaction of α_2_δ’s with other proteins (for review see Dolphin, 2018a). For example, we have demonstrated an interaction between α_2_δ-1 and Low-Density Lipoprotein (LDL) Receptor-related Protein-1 (LRP1), a multifunctional receptor which mediates cargo trafficking. LRP1 and its chaperone Rap, promote α_2_δ-1 maturation and trafficking to the cell-surface, and also enhance Ca_V_2.2 currents (Kadurin et al., 2017). Since LRP1 has multiple intracellular and extracellular protein ligands, and contains several cytoplasmic adaptor protein binding motifs (Lillis et al., 2008), it may play a role in α_2_δ-1 trafficking via recycling endosomes.

In another study, neurexin-1α enhanced Ca_V_2.1 currents when expressed with α_2_δ-1 but had no effect in the presence of α_2_δ-3 (Brockhaus et al., 2018), although no selective and specific interaction with neurexin-1α could be demonstrated, in contrast to a previous study (Tong et al., 2017). α_2_δ-1 has also been reported to enhance synaptogenesis, independently of Ca_V_ channels, through a gabapentin-sensitive extracellular interaction between the VWA domain of α_2_δ-1 and the Epidermal Growth Factor (EGF) domain of Thrombospondins (TSPs) 1-4, which are extracellular matrix proteins (Eroglu et al., 2009). More recently, another study (Lana et al., 2016) showed a weak interaction between α_2_δ-1 and TSP4, but was unable to demonstrate any interaction between cell-surface expressed α_2_δ-1 and TSP4 nor were they able to demonstrate the importance of the EGF domains of TSP4 in this interaction, concluding that α_2_δ-1 and TSP4 may only interact intracellularly at high concentrations. A further study showed a low affinity interaction between α_2_δ-1 and TSP4, but not other TSPs, which was gabapentin-insensitive and did not involve the EGF domain of TSP4 (El-Awaad et al., 2019). However, any interaction of α_2_δ subunits with TSPs is unlikely to be involved in their Rab11-dependent recycling.

A number of studies have also linked Rab11a function to growth cone development (Goldenring, 2015; Homma and Fukuda, 2016; van Bergeijk et al., 2015). Since we have demonstrated distinct trafficking pathways for α_2_δ-1 and α_2_δ-2 compared to α_2_δ-3 in relation to Rab11a-dependent recycling, it is possible that other demonstrated subtype-specific functions of α_2_δ proteins (Geisler et al., 2019; Kurshan et al., 2009) may also involve distinct trafficking pathways.

## Materials and Methods

### Molecular biology

The following cDNAs were used: Ca_V_2.2 (rabbit, D14157), Ca_V_2.2-HA and Ca_V_2.2-BBS (Cassidy et al., 2014), GFP-Ca_V_2.2-HA (Macabuag and Dolphin, 2015), β1b (rat, X61394) (Pragnell et al., 1991), α_2_δ-1 (rat, M86621) (Kim et al., 1992), α_2_δ-1-HA (Davies et al., 2006; Kadurin et al., 2012), α_2_δ-1-BBS, in which the BBS tag from α_2_δ-2 (Tran-Van-Minh and Dolphin, 2010) was inserted into α_2_δ-1 in the same position as the HA tag, α_2_δ-3 (AJ010949), α_2_δ-3-HA (Kadurin et al., 2016). mCherry (Shaner et al., 2004) or CD8 (Rougier et al., 2005) were included as a transfection markers where stated. Human Rab11a (AF000231) containing the S25N mutation (Dupre et al., 2006; Tran-Van-Minh and Dolphin, 2010), was obtained from From Prof. T. Hébert, McGill University.

### Antibodies and other materials

Antibodies (Abs) used were: anti-α_2_δ-1 Ab (mouse monoclonal, Sigma-Aldrich), anti-α_2_δ-3 Ab (Davies et al., 2010), anti-HA Ab (rat monoclonal, Roche), anti-Ca_V_2.2 II-III loop Ab (rabbit polyclonal) (Raghib et al., 2001). The following secondary Abs (raised in goat) were used: anti-rat or anti-rabbit IgG-Alexa Fluor (AF) 488 or 594 (ThermoFisher), biotin-labelled anti-rat IgG (Sigma). Other reagents used were BTX-AF488 (Invitrogen) and streptavidin-AF633 (Invitrogen).

### Cell lines: culture and transfection

tsA-201 cells were grown in Dulbecco’s Modified Eagle Medium (DMEM) (Invitrogen) supplemented with 10 % Foetal Bovine Serum, 50 U/ml penicillin/streptomycin and 1 % GlutaMAX (Invitrogen). Cells were maintained in tissue culture flasks at 37°C in a humidified atmosphere of 5 % CO_2_ and grown up to 70-90 % confluence prior to transfection or further passage. Neuro2a (N2a) cells were cultured in 50 % DMEM (with high glucose and L-glutamine) and 50 % OPTI-MEM (with L-glutamine), supplemented with 50 U/ml penicillin/streptomycin, 5 % FBS, and 1 % GlutaMAX (Life Technologies). The N2a cells were cultured to 80 % confluence in a 5 % CO_2_ incubator at 37 °C and passaged every 3 to 4 d.

For transient expression, N2a or tsA-201 cells were transfected using either PolyJet™ (SignaGen Laboratories) or Fugene® 6 (Promega) in a 3:1 ratio with cDNA mix, according to the manufacturers’ instructions. A typical cDNA mix was 2μg total DNA per 35 mm culture dish at a ratio of 3:2:2 of Ca_V_2.2: β1b: α_2_δ. In experiments using Rab11a(S25N) the cDNA mix was 3:2:2:1 for Ca_V_2.2: β1b: α_2_δ: Rab11a(S25N). In conditions lacking a given construct, the DNA mix was supplemented with the appropriate empty vector to have the same total DNA. For fluorescence imaging experiments in either tsA-201 or N2a cells, all cDNAs were in a pcDNA3, pRK5 or pCMV vector. In electrophysiological experiments using tsA-201 cells, all cDNAs were in a pMT2 vector, with the exception of Rab11a(S25N) (pCMV).

### Hippocampal neuronal culture and transfection

Hippocampal neurons were isolated from the hippocampus of P0 rats, which were killed under a UK Home Office Schedule 1 procedure. The cerebrum was cut into two, and the hippocampi were extracted from each hemisphere in ice cold HBSS with 10 mM HEPES. The hippocampi were then cut into small segments and digested gently in Papain solution (7 U/ml Papain, 0.2 mg/ml L-Cysteine, 0.2 mg/ml Bovine Serum Albumin (BSA), 5 mg/ml glucose, 10 mM HEPES in HBSS with 30 Kunitzs/ml DNase) in a shaking water both for 40 min at 37 °C. The cells were washed twice with the growth medium (NeuroBasal, 2 % B27supplement, 1 unit/ml penicillin, 1 μg/ml streptomycin, 1 % Glutamax, 0.1% β-mercaptoethanol), and triturated gently. The cells were plated onto the coverslips that are coated with poly-L-lysine and laminin at 750 cells/μl, 100 μl per coverslip. The entire growth medium was changed after 2 h of plating, and half of the medium was then changed every 3-4 d.

Hippocampal neurons were transfected after 7 d in culture, using cDNAs cloned into pCAGGS. 2 h prior to transfection, the growth medium was replaced with a mixture of 50 % conditioned, and 50 % fresh growth medium. The transfection process was as follows: 4 μg cDNA was mixed in a 50 μl total with OPTI-MEM™ (Thermo Fisher), 2 μl Lipofectamine®2000 (Thermo Fisher) was mixed in a 50μl total volume of OPTI-MEM™ in separate tubes by pipetting; mixing was done by gentle pipetting. DNA and transfection reagent mixes were then combined and incubated for 5 min at room temperature before being added dropwise to cultures. Hippocampal neurons were cultured for a further 7 d after transfection before fixation and immunostaining.

### Immunocytochemistry

Cells were plated onto either coverslips or glass-bottomed dishes (MatTek Corporation) which were coated with poly-L-lysine prior to transfection, and cultured in a 5 % CO_2_ incubator at 37°C. After 36 - 48 h expression, cells were fixed with ice cold 4 % paraformaldehyde (PFA), 4 % sucrose in PBS, pH 7.4 at room temperature for 5 min. Where antigen retrieval was used, fixed samples were incubated in pH 6 citrate buffer at 98°C for 10 min (Cassidy et al., 2014). For labelling the HA epitope on the cell-surface in non-permeabilised conditions, cells were incubated with primary Ab with 2 % BSA and 10 % goat serum in PBS at room temperature for 1 h for cell lines or overnight at 4°C for neurons. When permeabilization was performed, cells were incubated with 0.2% Triton X-100 in PBS for 5 min (Cassidy et al., 2014). The secondary Ab was added with 2.5 % BSA and 10 % goat serum in PBS and incubated for 1 h at room temperature. Cell nuclei were stained with 0.5 µM 4’,6’-diamidino-2-phenylindole (DAPI) in PBS for 10 min. The coverslips were mounted onto glass slides using VECTASHIELD® mounting medium (Vector Laboratories). In experiments probing for both cell-surface and intracellular HA epitopes (Fig. 1), cells were fixed and blocked as described above, then immunostained for 1 h with rat anti-HA Ab (1:500), followed by 1 h incubation with biotin-labelled anti-rat IgG (1:1000). Following this, cells were permeabilised with 0.2 % Triton-X100 for 5 min and re-probed with rat anti-HA Ab for 1 h. Cells were then incubated with anti-rat-AF594 Ab (1:500) and streptavidin-AF633 (1:500), to visualise intracellular and extracellular HA epitopes, respectively. In experiments probing for cell surface BBS tag in Ca_V_2.2 (Fig. 2, Fig. S1), cells were live-labelled with BTX-AF488 (10 µg/ml, Invitrogen) for 30 min at 17 oC. Cells were then washed, fixed, permeabilised and immunostained, using a rabbit Ab against the II-III loop to detect total Ca_V_2.2, followed by secondary anti-rabbit-AF594 Ab.

### Net forward trafficking assay

Transfected cells were plated onto 22 x 22 mm glass coverslips coated with poly-L-lysine, and cultured in a 5 % CO_2_ incubator at 37°C. After 40 h expression, N2a or tsA-201 cells were washed twice with Krebs-Ringer-HEPES (KRH) buffer and incubated with 10 μg/ml unlabelled BTX (Life technologies) for 30 min at 17°C. The unbound BTX was washed off with KRH, and the cells were then incubated with BTX-AF488 (10 µg/ml) in KRH at 37°C. To terminate the reaction, cells were washed twice with cold KRH and then fixed with 4 % PFA, 4 % sucrose in PBS at specified times for the kinetic assay. After fixation, cells were permeabilised and intracellular expression markers and/or nuclei were labelled as described above. The coverslips were then mounted onto glass slides using VECTASHIELD® mounting medium.

### Confocal microscopy

All images were acquired using an LSM 780 scanning confocal microscope (Zeiss), equipped with a Plan-Apochromat 63x/1.4 or 40x/1.3 DICII oil immersion objective lens, in16-bit mode. For each experiment, the laser power, gain and acquisition settings were kept constant between images that were used for quantification, although laser power and gain settings were adjusted between experiments depending on expression and staining level of the samples. Where possible, the region of interest was determined by identifying cells with expression of a transfection marker or intracellular staining of the protein of interest (e.g. GFP, HA), without selecting for the cell-surface immunostaining, to avoid bias. In experiments where an appropriate intracellular expression marker was absent, nuclear staining with DAPI was used to identify viable cells (having an intact nucleus); cell-surface measurements were made for all viable cells per field of view. In addition, a 2×2 or 3×3 tile scan was performed to further remove the bias in selecting cells with high expression. For cell-surface expression analysis, images were taken usually with 1 μm optical section. Confocal images were imported and analysed in ImageJ (National Institutes of Health). The plasma membrane fluorescence was quantified using the freehand brush tool with a selection width of 0.66 μm and tracing the membrane region manually. Total or intracellular fluorescence was quantified using the freehand selection tool, omitting the signal intensity from the nuclei. The background fluorescence in each channel was measured and subtracted from mean cell-surface or intracellular fluorescence measurements in image analysis.

For analysis of neurite expression in hippocampal neurons, an average of 10 - 15 cells were selected per condition for an individual experiment. Neurons were selected based on expression of free mCherry as a transfection marker. Between 2 - 5 neurites were measured per mCherry-positive cell to reduce over-representation of highly branched neurons. The freehand brush tool was used to manually trace neurites of lengths between 25 - 60 µm, and width of 2 µm, beginning at a distance of 100 µm from the cell soma. Neurites were selected and traced using mCherry to avoid bias, measurements were then taken in channels for expression of HA and/or GFP tags on expressed Ca_V_2.2 constructs. Background fluorescence measurements were taken for each condition and subtracted from mean fluorescence values during image analysis. Mean neurite signal intensity for each channel was calculated for neurons of a given condition and normalized to the stated controls. Normalized data were then pooled between experiments with a minimum of three separate experiments.

In cell-surface expression and forward trafficking experiments, where relevant, all cells chosen for analysis contained intracellular Ca_V_2.2 expression. For the forward trafficking assays, the membrane fluorescent intensities were fitted to the single exponential association. All experiments were repeated n = 3 - 5 times, and approximately 30 - 50 cells were analysed for each experiment. All data are presented as pooled cells, except for the trafficking initial rates, which are averages of separate experiments.

### Electrophysiology

tsA-201 cells were transfected with cDNA mix containing Ca_V_2.2, α_2_δ-1, β1b, and CD8 at a ratio of 3:2:2:0.8 using a Fugene transfection protocol. After 40 h expression, cells were replated in cell culture medium at 1 in 3 or 1 in 5 dilution depending on their confluency. Transfected cells were identified by CD8 expression was detected with CD8 Dynabeads (Life Technologies). Whole-cell currents were recorded in voltage-clamp mode in the following solutions; intracellular solution (mM): 140 Cs-aspartate, 5 EGTA, 2 MgCl_2_, 0.1 CaCl_2_, 2 K_2_ATP, 20 HEPES, pH 7.2, 310 mOsm; extracellular solution (mM): 1 BaCl_2_, 3 KCl, 1 NaHCO_3_, 1 MgCl_2_, 10 HEPES, 4 D-glucose, 160 tetraethylammonium bromide, pH 7.4, 320 mOsm. The borosilicate glass electrode resistance was between 1.5 and 4 MΩ. Cell capacitance and series resistance were compensated to 60 - 70 %. Whole-cell currents were recorded on Axopatch-200B amplifier using pClamp 9 or 10 (Molecular Devices). The cells were held at −90 mV, and 50 ms pulses were applied in +10 mV steps between − 50 mV and +50 mV. To correct for the leak current, P/8 leak subtraction protocol was applied. Recordings were made at 20 kHz sampling frequency and filtered at 5 kHz (lowpass 4-pole Bessel filter) in the amplifier. The digital low-pass 8-pole Bessel filter with 1 kHz 3dB cut-off was applied in Clampfit 10.7 (Molecular Devices) before the current amplitudes were determined. Average peak currents were taken between 8 - 13 ms after the test potentials were applied and normalized to the cell capacitance to obtain current density. *IV* relationships were fit by a modified Boltzmann equation as follows: I = *G*_*max*_*\*(V-V*_*rev*_*)/(1+exp(-(V-V*_*50, act*_*)/k)*), where I is the current density (in pA/pF), *G* _*max*_ is the maximum conductance (in nS/pF),*V*_*rev*_ is the apparent reversal potential (mV), V_50, act_ is the midpoint voltage for current activation (mV), and *k* is the slope factor.

## Statistical analysis

The number of samples (n) in most of the experiments in this study is the total number of individual cells analysed, which are pooled from a minimum of three separate experiments unless stated otherwise. Data from individual experiments were normalized to their control conditions prior to being pooled to account for inter-experimental variance. Mean values and standard error of mean (SEM) were calculated for normalized, pooled data. The only exception is for the experiments determining the rates of Ca_V_2.2 trafficking, where the mean and SEM values were determined from the mean of repeated experiments. Statistical analysis was performed using Student’s t test or one-way ANOVA with appropriate post-hoc test as stated, in GraphPad Prism 7.

## Acknowledgements

The authors thank Wendy S Pratt and Kanchan Chaggar for technical support, and Dr. Laurent Ferron and Kjara Pilch for hippocampal cultures. We thank Drs. Karen Page and Manuela Nieto-Rostro for commenting on the manuscript.

## Funding

This research was funded, in part, by the Wellcome Investigator award to ACD (098360/Z/12/Z)]. For the purpose of Open Access, the corresponding author has applied a CC BY public copyright licence to any Author Accepted Manuscript version arising from this submission. JOM was supported in part by the UCL Grand Challenges PhD program.

## Competing interests

The authors declare no competing interests

## Author Contributions

JOM performed all experimental work, and analysed data. JOM and ACD conceived the study and co-wrote the manuscript.

## Supplementary Figures

**Fig. S1.**
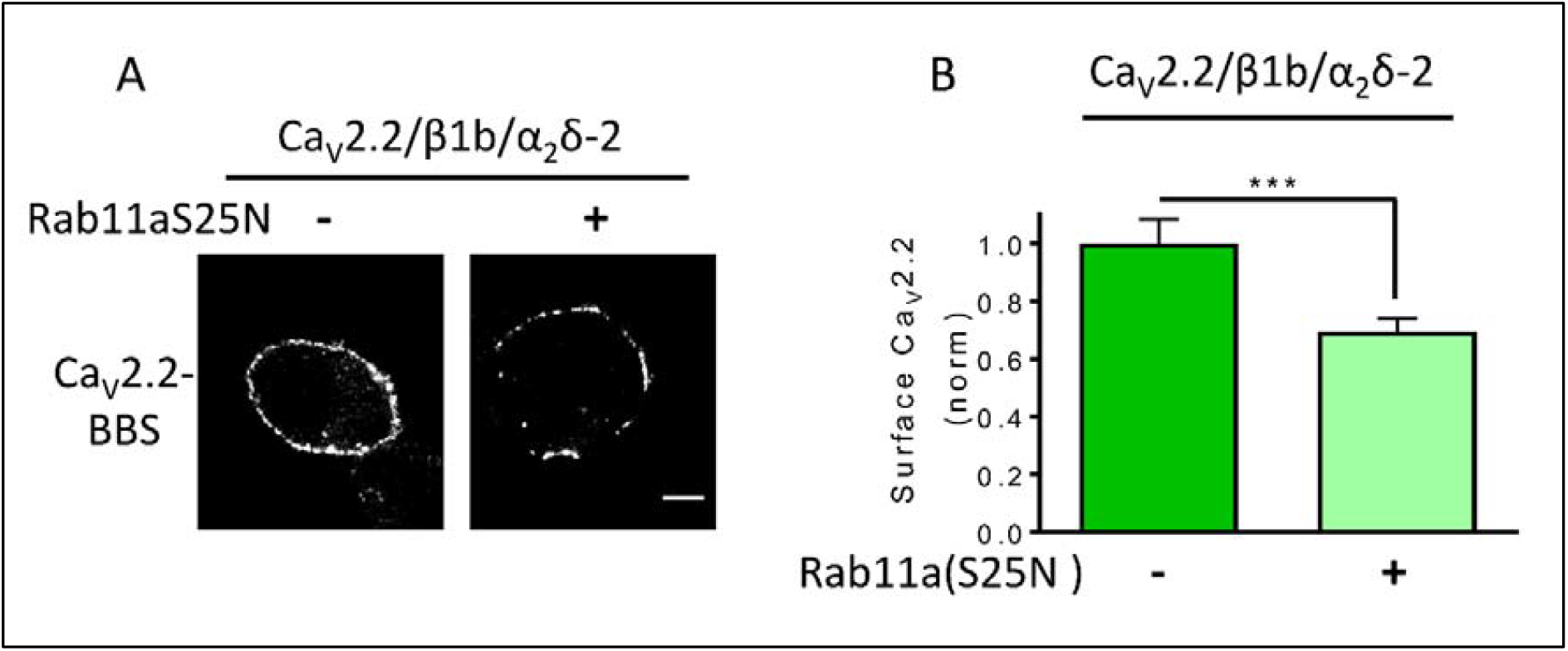
Steady-state cell-surface Ca_V_2.2 is reduced by Rab11a(S25N) when expressed with α_2_δ-2. **(A)** Confocal images of cell-surface Ca_V_2.2-BBS/β1b expressed in N2a cells with α_2_δ-2 in the absence or presence of Rab11a(S25N). Scale bar = 5 µm. **(B)** Normalized mean cell-surface Ca_V_2.2-BBS with β1b and either: α_2_δ-2 (green, n = 97 cells), α_2_δ-2 + Rab11a(S25N) (light green, n = 114 cells). Mean fluorescence intensity per cell was normalized to Ca_V_2.2/β1b/α_2_δ-2 controls, pooled from three separate transfections, and data are plotted as mean ± SEM values. Statistical significance was determined using Student’s unpaired t test, \*\*\**P* <0.001.

**Fig. S2.**
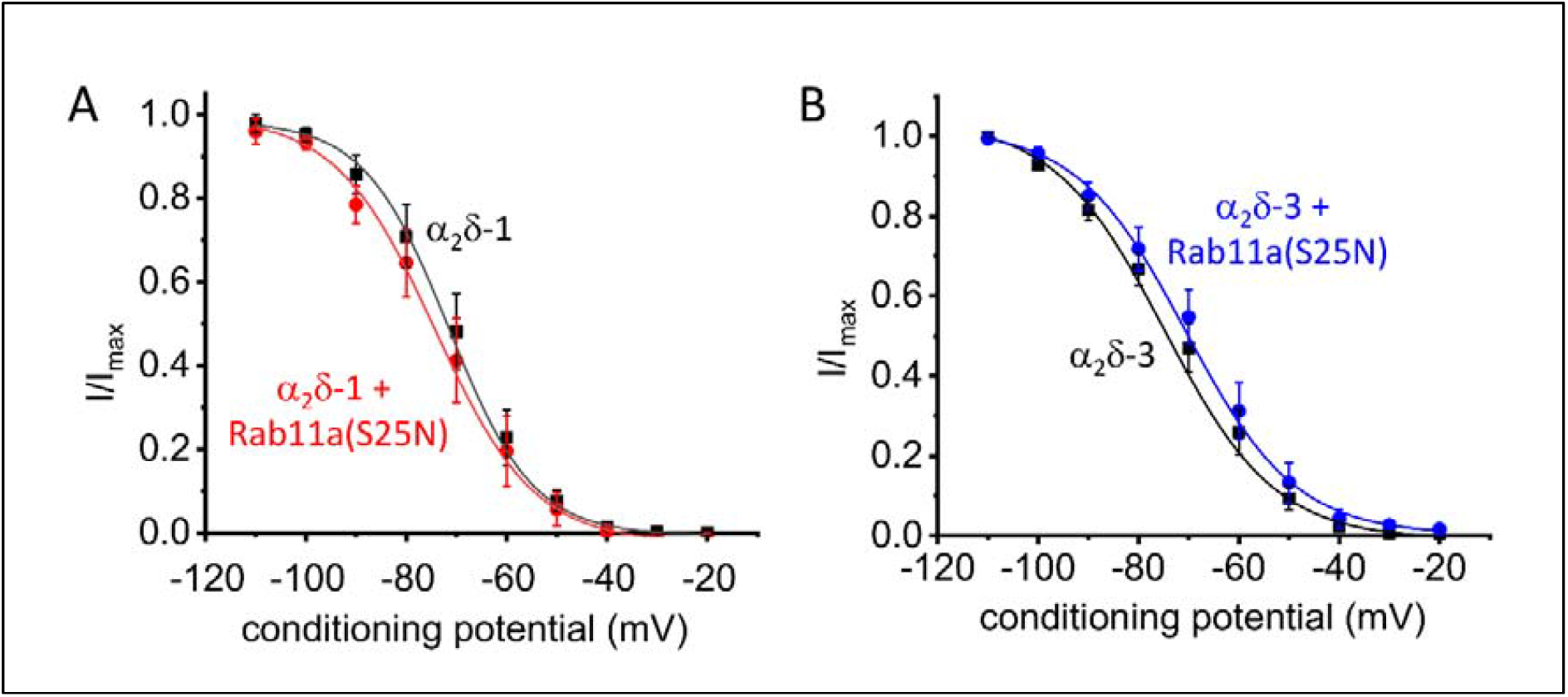
Rab11a(S25N) has no effect on steady-state inactivation of whole-cell Ca_V_2.2 currents when expressed with α_2_δ-1 or α_2_δ-3. **(A)** Mean steady-state inactivation for Ca_V_2.2 with: β1b/α_2_δ-1 (black squares; n = 7 cells) or β1b/α_2_δ- 1 + Rab11a(S25N) (red circles; n = 7 cells), fit to a Boltzmann function. The V_50, inact_ was −71.8 ± 3.6 mV and −73.8 ± 3.9 mV, respectively. **(C)** Mean steady-state inactivation for Ca_V_2.2 with: β1b/α_2_δ-3 (black squares; n = 7 cells) or β1b/α_2_δ- 3 + Rab11a(S25N) (blue circles; n = 7 cells), fit to a Boltzmann function, The V_50, inact_ was −71.5 ± 2.5 mV and −69.3 ± 3.4 mV, respectively.

## Notes

### Competing Interest Statement

The authors have declared no competing interest.

